# Effects of antiretroviral treatment on central and peripheral immune response in mice with EcoHIV infection

**DOI:** 10.1101/2024.04.11.589109

**Authors:** Qiaowei Xie, Mark D Namba, Lauren A Buck, Kyewon Park, Joshua G Jackson, Jacqueline M Barker

## Abstract

HIV infection is an ongoing global health issue despite increased access to antiretroviral therapy (ART). People living with HIV (PLWH) who are virally suppressed through ART still experience negative health outcomes, including neurocognitive impairment. It is increasingly evident that ART may act independently or in combination with HIV infection to alter immune state, though this is difficult to disentangle in the clinical population. Thus, these experiments used multiplexed chemokine/cytokine arrays to assess peripheral (plasma) and brain (nucleus accumbens; NAc) expression of immune targets in the presence and absence of ART treatment in the EcoHIV mouse model. The findings identify effects of EcoHIV infection and of treatment with bictegravir (B), emtricitabine (F) and tenofovir alafenamide (TAF) on expression of numerous immune targets. In the NAc, this included EcoHIV-induced increases in IL-1α and IL-13 expression and B/F/TAF-induced reductions in KC/CXCL1. In the periphery, EcoHIV suppressed IL-6 and LIF expression, while B/F/TAF reduced IL-12p40 expression. In absence of ART, IBA-1 expression was negatively correlated with CX3CL1 expression in the NAc of EcoHIV-infected mice. These findings identify distinct effects of ART and EcoHIV infection on peripheral and central immune factors and emphasize the need to consider ART effects on neural and immune outcomes.

## Introduction

Worldwide, there are approximately 39 million people living with HIV (PLWH), 75% of whom are accessing antiretroviral therapy (ART) (UNAIDS, 2023). HIV infection is associated with immune responses that include increased circulating cytokine levels in the periphery and within the central nervous system (CNS), and which may contribute to severity of disease (Merrill and Chen, 1991). Although ART can effectively maintain HIV-1 viral load at undetectable levels and prolong lifespan of PLWH, ART does not completely eliminate infection or the impact of HIV-induced immune response peripherally or within the central nervous system (CNS) on comorbidities including HIV-associated neurocognitive disorder (HAND) (Valcour et al., 2012; Ripamonti and Clerici, 2021; Guha et al., 2023).

HIV-1 enters the CNS via infected peripheral monocytes and T cells which can cross the blood brain barrier (Alexaki et al., 2008; Williams et al., 2014; Veenstra et al., 2017a). Once within the CNS, HIV can infect glial cells including microglia and astrocytes - but not neurons-, establishing a viral reservoir that persists despite ART treatment (Hauser et al., 2007; Eugenin et al., 2011; Wallet et al., 2019). The presence of HIV and HIV proteins in the brain can induce release of inflammatory factors, contributing to neuroinflammation and neuronal dysfunction (Potter et al., 2013; Hauser and Knapp, 2014). These deficits may be further exacerbated within reward-related substrates as findings from both clinical populations and preclinical models identify deficits within the nucleus accumbens (NAc) and its connecting structures that may ultimately contribute to aberrant reward processing, learning, and memory seen in PLWH (Paul et al., 2005; Roscoe et al., 2014; McLaurin et al., 2018; Ezeomah et al., 2022). For example, HIV dampens dopamine (DA) system function and dysregulates glutamatergic synaptic plasticity in the NAc, likely through perturbations in immune signaling (Buch et al., 2011; Illenberger et al., 2020). Considering the high rates of comorbid substance use disorder (SUD) and HIV infection, investigating neuroimmune changes in the NAc may provide insight into the development of novel approaches to mitigate addiction-related behaviors and neurocognitive outcomes in PLWH with SUD.

Beyond the persistence of inflammatory response in PLWH even in the presence of ART, it is becoming increasingly evident that ART interacts with HIV and/or acts independently to produce a distinct immune profile in the periphery and CNS (Hileman and Funderburg, 2017; Hughes et al., 2020; Bertrand et al., 2021; Yuan and Kaul, 2021; Nguyen et al., 2023). In clinical findings, the direct effects of ART are difficult to disentangle from interactions with aging, viral status, and additional comorbidities. However, preclinical models point to ART-induced dysregulation of peripheral and central nervous system immune function (Zulu et al., 2021). Cognitive outcomes include spatial learning and memory impairments following treatment with nevirapine (Zulu et al., 2020) or emtricitabine + tenofovir disoproxil fumarate (Zulu et al., 2021) in the absence of infection. Thus, preclinical research will be essential to fully characterize the independent and interactive effects of ART on neural and immune outcomes.

Together, these findings point to a need to understand the independent and interactive effects of HIV infection and ART on peripheral and CNS immune outcomes, necessitating preclinical models of HIV (Buch et al., 2011; Gorantla et al., 2012; Edagwa et al., 2014; Namba et al., 2023a). To accomplish this, the current study utilized the EcoHIV infection mouse model, in which a chimeric virus is generated by replacing the coding region of the HIV-1 glycoprotein gp120 with that of gp80 (Potash et al., 2005). Here, infection was initiated in the periphery. To model a commonly prescribed ART regimen, the current study used a daily combination treatment with bictegravir (B) – a recommended integrase strand transfer inhibitor (INSTI) -, emtricitabine (F) and tenofovir alafenamide (TAF) – nucleoside reverse transcriptase inhibitors (NRTIs) -, in accordance with recommended initial ART regimens that show high rates of viral suppression in PLWH (Havens et al., 2023; Sax et al., 2023).

Experiments in this study assessed immune factors in plasma and in the NAc, a key substrate of brain reward circuitry known to be dysregulated in models of HIV (McLaurin et al., 2018; Namba et al., 2023b), including following CNS infection with EcoHIV (Li et al., 2021). Neuroimmune dysregulation in this region can impair excitatory glutamatergic neuronal plasticity, thus driving increased susceptibility to addictive drugs (Lacagnina et al., 2017; Namba et al., 2021). Given the evidence that microglia are the principle HIV-1 reservoir within the CNS and the known role of microglia in sensing neural activity and modulating neuronal plasticity (Wallet et al., 2019), we investigated EcoHIV and ART effects on microglia. Fractalkine (CX3CL1) is a soluble chemokine released by neurons, it is an exclusive ligand for CX3CR1 receptors that are expressed on microglia (Chapman et al., 2000). CX3CL1/ CX3CR1 signaling has been proposed to mediate neuron-microglia communication that underlies microglial activation and neuroimmune response (Mizuno et al., 2003; Pawelec et al., 2020) and thus we further characterized NAc expression of Iba-1 and CX3CL1. The present results identify independent and interactive effects of both EcoHIV infection and B/F/TAF treatment on cytokine expression in both plasma and the NAc.

## Materials and Methods

### Subjects

Adult male (n=24) and female (n=24) C57BL/6J mice (9 weeks upon arrival) were obtained from Jackson Laboratories. Following arrival, mice were group housed in same-sex cages for 7 days to acclimatize with *ad libitum* access to a standard chow diet and water. Mice were housed at the Drexel University College of Medicine under standard 12-hour light:12-hour dark conditions in microisolation conditions. All experiments were approved by the Institutional Animal Use and Care Committee at Drexel University.

### EcoHIV virus generation

Plasmid DNA encoding the EcoHIV-NDK coding sequence (gift from Dr. David Volsky) was purified from bacterial stocks (Stbl2 cells, ThermoFisher#10268019) and purified using an endotoxin free plasmid purification kit (ZymoPure #D4200). Purified DNA was transfected into nearly confluent (80-90%) 10 cm^2^ plates of low passage LentiX 293T (Takara #632180) using a calcium phosphate transfection protocol. The cell culture supernatant was collected at 48 hours post-transfection and cellular debris was pelleted by centrifugation at low speed (1500 x g) on a benchtop centrifuge (4°C) followed by passage through a cell strainer (40µm). Supernatant containing viral particles was mixed 4:1 with a homemade lentiviral concentrator solution (4x; MD Anderson) comprised of 40% (w/v) PEG-8000, 1.2M NaCl in PBS (pH 7.4). The supernatant-PEG mixture was placed on an orbital shaker (60 rpm) and incubated overnight at 4°C. This mixture was centrifuged at 1500 x g for 30 minutes at 4°C. After centrifugation, the medium was removed, and the pellet rinsed 1x with PBS. The pellet was centrifuged again (1500 x g for 5 min), solution removed, and the viral pellet resuspended in cold, sterile PBS. Viral titer (p24 core antigen content) was determined initially using a LentiX GoStix Plus titration kit (Takara #631280) and subsequently by HIV p24 AlphaLISA detection kit (PerkinElmer # AL291C). Viral stocks were aliquoted and stored at -80°C until used.

### EcoHIV inoculation and antiretroviral drug self-administration

Following 1 week of acclimation, mice were matched based on sex into EcoHIV or sham control groups. Mice were inoculated with EcoHIV at a dose of 300ng p24 EcoHIV-NDK, i.p. For inoculation, mice were lightly anesthetized with 5% isoflurane and injected interperitoneally with either EcoHIV or vehicle (sterile 1x PBS) for sham controls and subsequently individually housed.

Antiretroviral drug powder (bictegravir, emtricitabine, and tenofovir alafenamide, B/F/TAF, Cayman Chemicals), was pre-mixed with Rodent Liquid Diet (Cat#F1268, AIN-76, Bio-Serv, NJ, USA) as a liquid mixture. Mice were assigned based on sex and infection status into B/F/TAF-treated and vehicle-treated groups. Beginning one week after inoculation, EcoHIV-infected and sham control mice were single-housed for control of daily B/F/TAF dosing via oral self-administration. Mice consumed 5 mL B/F/TAF mixture, dosed as 0.375 mg B/1.23 mg F/0.15375 mg TAF per mouse diet per day (calculated as the human equivalent dose (Nair and Jacob, 2016; Daar et al., 2018; Molina et al., 2018)), or received 5 mL liquid diet as vehicle controls daily until the end of the test. To ensure controlled dosage consumption, mice were subjected to food restriction, and additional food pellet increments were administered as needed to maintain body weights at pre-restriction levels.

### Brain and blood sample collection and preparation

#### Blood samples

Blood samples were collected from all mice 1, 3 and 5 weeks following EcoHIV inoculation. For blood collections, mice were briefly anesthetized with 5% vaporized isoflurane and, using the submandibular collection method, blood was collected from each mouse. Blood samples were centrifuged at 8700 x g for 20 minutes. Plasma was collected from supernatant and stored at -80°C. Following bleed collection at the end of week 5, mice were overdosed on isoflurane and euthanized via rapid decapitation, brains and spleen were removed, flash-frozen on dry ice, and stored at -80°C.

#### Nucleus accumbens (NAc) lysate

Frozen brains were sectioned and bilateral NAc was isolated using a biopsy punch and collected into 1.5 mL tubes. NAc brain tissues were homogenized using 1% SDS in RIPA buffer and incubated at 4°C for 30 mins. Homogenized NAc samples were centrifuged at 12,000 x g for 10 mins. The supernatants were collected for whole tissue lysate. Protein concentration was quantified using Pierce™ BCA Protein Assay Kit (Thermo Scientific, Waltham, MA, USA).

### EcoHIV Infection Validation

Spleens were harvested at the time of euthanasia and flash frozen for storage at -80°C. DNA was isolated from spleens using the Qiagen QIAamp DNA Mini Kit (#51304). Confirmation of infection was performed by the University of Pennsylvania Center for AIDS Research (CFAR). The CFAR received DNA samples for cell-associated DNA assay and performed qRT-PCR using HIV-LTR primers and probe provided below. The OD was used to estimate the input cell numbers to normalize the data.

Kumar LTR F, GCCTCAATAAAGCTTGCCTTGA

Kumar LTR R, GGGCGCCACTGCTAGAGA

Kumar LTR Probe (FAM/BHQ), 5’ CCAGAGTCACACAACAGACGGGCACA 3’

### Quantification of NAc and Peripheral Immune Proteins by Mouse Cytokine 32-PlexDiscovery Assay

Cytokine and chemokine expression in NAc lysates and plasma were analyzed by multiplex immunoassay. Mouse NAc lysates were diluted 1:1 in PBS (PH∼7.5) prior to shipping. Plasma was sent undiluted. Mouse cytokines, chemokines and growth factors in NAc lysate and plasma were measured using the Mouse Cytokine 32-PlexDiscovery Assay (Eve Technologies, MD32, Calgary, AB, Canada). The multiplex analysis was performed using the Luminex 200 system (Luminex, Austin, Texas) by Eve Technologies Corporation. Results that were out of range (i.e., outside the standard curve) were excluded from the dataset prior to analysis. The assay sensitivities of the mouse markers range from 0.3-30.6 pg/mL. The 32 targets included: Eotaxin, G-CSF, GM-CSF, IFNγ, IL-1α, IL-1β, IL-2, IL-3, IL-4, IL-5, IL-6, IL-7, IL-9, IL-10, IL-12p40, IL-12p70, IL-13, IL-15, IL-17A, IP-10, KC, LIF, LIX, MCP-1, M-CSF, MIG, MIP-1α, MIP-1β, MIP-2, RANTES, TNFα, VEGF.

### Western blot

Aliquots of NAc lysate from the same samples used for the cytokine assay were analyzed for Iba-1 and CX3CL1 expression via western blot. The NAc tissue was lysed into whole tissue lysate in RIPA buffer containing protease inhibitor, clarified by centrifugation, and the protein concentrations were determined using a BCA Protein Assay. Equal amounts of protein (12 μg) were loaded and run on 4-12% gradient gel (NuPAGE^TM^, 4-12%, Bis-Tris, 1.0 mm). Proteins were transferred onto nitrocellulose membrane using an iBlot^TM^ 2 Gel Transfer Device. The membranes were then blocked using 5% BSA (Iba-1) or 5% milk (CX3CL1) in Tris-buffered saline + 0.1% Tween-20 (TBS-T) for 2 hours. Rabbit anti-Iba-1 antibody (FUJIFILM Wako Pure Chemical Corporation, 019-19741; 1:500), rabbit anti-CX3CL1 (Bioss, bs-0811R; 1:500), or rabbit anti-GAPDH antibody (1:3000) was added to the blocking buffer, and membranes were incubated overnight at 4°C. The membranes were then washed in TBS-T and incubated with horseradish peroxidase-labeled (HRP) goat anti-rabbit secondary antibody (Abcam ab97080; 1:3000 for Iba-1 and CX3CL1, 1:10,000 for GAPDH) for 1 hour. Chemiluminescent substrate (SuperSignal™ West Pico Plus, Thermo Fisher) was used to develop the membranes for protein detection. Protein bands were analyzed using ImageJ. A between-gel control sample was loaded onto every gel and the expression of Iba1 and CX3CL1 was normalized to this sample prior to normalizing to GAPDH to control for between-gel differences. Thus, data were calculated as ratios of normalized Iba-1 or CX3CL1/normalized GAPDH, and then as a fold change relative to the mean of control group (vevehicle mice).

### Statistical Analyses

All data were analyzed in GraphPad Prism. Body weights were analyzed by three-way mixed model ANOVA using Greenhouse-Geisser correction. Cytokine/chemokine data were analyzed by two-way ANOVA or by unpaired t-test. Analyses were log2 transformed or square root transformed when normal distribution was not met. Significant interactions were followed by Tukey’s post hoc analysis. Correlational analyses were performed by simple linear regression. Significance levels for each test were to p < 0.05. Principal component analysis (PCA) was applied to identify common characteristics among the dependent variables using Prism. Raw expression data of cytokine/chemokine were scaled and centered. The correlation matrix and eigenvalue were calculated, eigenvalues greater than 1 were considered significant in contributing to the components, and data was plotted on the first two components. Z-score two population comparison was applied to compare the proportion of mice assigned to each quadrant following PCA.

## Results

### EcoHIV-NDK infection of wild type mice

Adult male and female C57BL/6J mice (9 weeks) were matched by sex to undergo either sham or EcoHIV inoculation and vehicle or B/F/TAF exposure (**Fig.1a**). To confirm EcoHIV-NDK infection status, terminal viral DNA burden was measured in spleens by qPCR. A subset of tissues (n = 7) was run from sham control mice to confirm that no viral burden was observed. EcoHIV-NDK-infected mice showed viral burdens in the range from 0.3-5 x 10^3^ viral DNA copies per 10^6^ spleen cells after 5 weeks of EcoHIV infection. One EcoHIV vehicle treated mouse was excluded from analyses as no viral DNA was detected. There was no significant difference in mean EcoHIV DNA viral burden between B/F/TAF treated mice (n = 12) and vehicle treated EcoHIV mice (n = 11) [t (21) =0.103, p = 0.9191; **Fig. 1b**]. Body weight was monitored 3 times per week following inoculation (**Fig. 1c**). No effects of EcoHIV infection [F (1, 20) = 1.325, p = 0.2633] or B/F/TAF treatment [F (1, 20) = 0.8443, p = 0.3691] were observed on change in weight. However, there was a main effect of time [three-way ANOVA; F (2.006, 40.12) = 37.13, p < 0.0001; Greenhouse-Geisser corrected, **Fig. 1c**]. A *post-hoc* Dunnett’s test revealed that weights were reduced on day 3 compared to day 0 (p<0.05). Weights were greater than day 0 on day 24, 27, 30 and 33 (p’s < 0.05).

**Figure 1.**
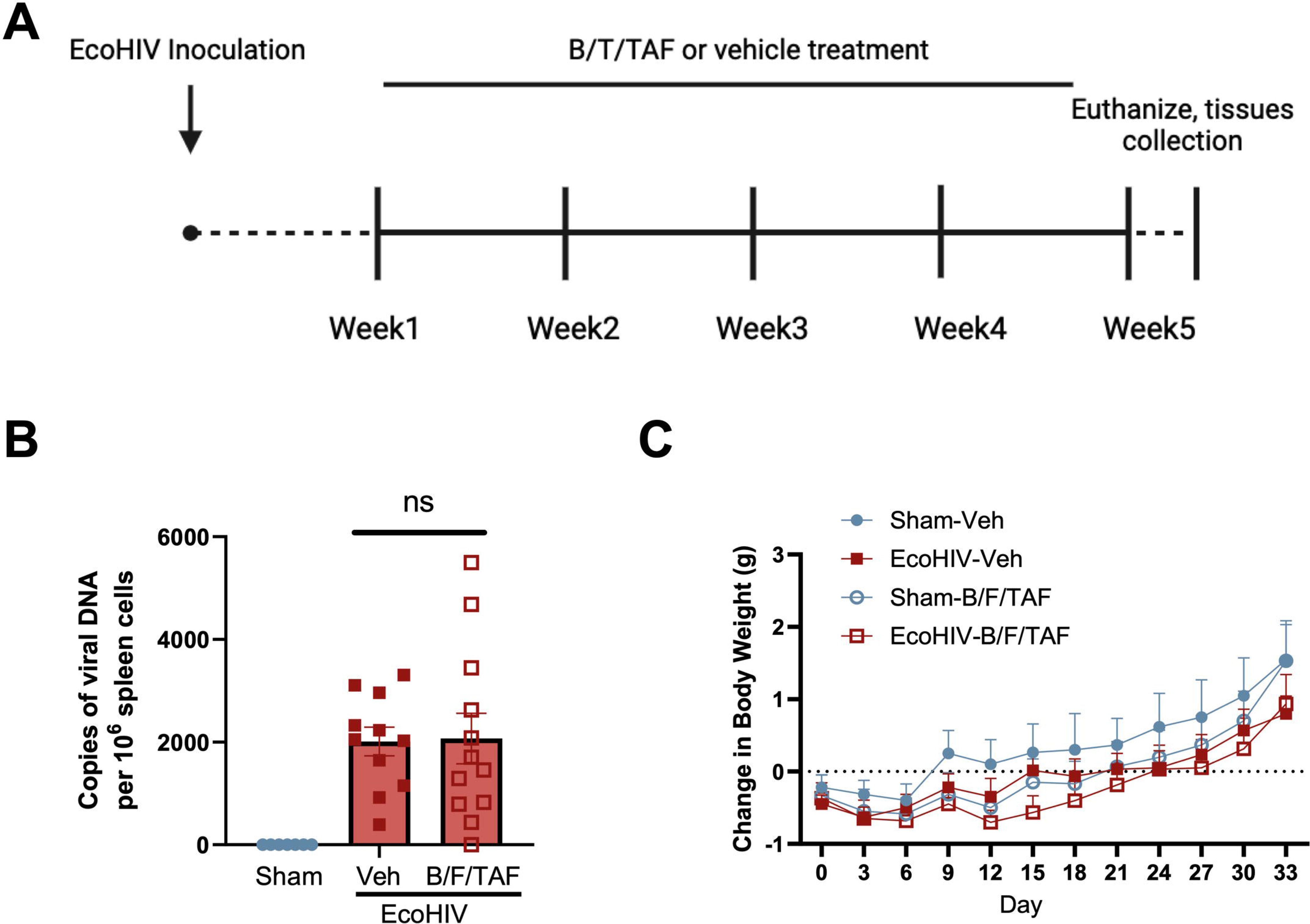
Mouse model of EcoHIV infection. (**a**) Timeline of experiments (created with Biorender.com). Mice were matched based on sex into EcoHIV or sham control groups and for treatment with B/F/TAF. (**b**) Copies of viral DNA euthanasia timepoint (week 5 of infection). The number of copies of viral DNA per 10^6^ spleen cells did not differ between EcoHIV-infected mice that were treated with vehicle (n=11) versus B/F/TAF (n=12). (**c**) Body weight changes verse baseline weight throughout 5 weeks of the experiment. No effects of EcoHIV infection or B/F/TAF treatment were observed on weight change. A main effect of day was observed such that weight increased across time. Circles and square represent sham and EcoHIV mice, open and closed symbols represent B/F/TAF and vehicle treated mice, respectively. Bars represent mean +/- SEM.

### EcoHIV infection alters neuroimmune responses in the NAc

To assess the expression levels of immune markers in the brain of EcoHIV infected mice, this study utilized mouse cytokine 32-PlexDiscovery Assay. Results for analytes that were outside the linear range of detection were excluded from the dataset prior to analysis. Two-way ANOVA revealed changes in immune markers. EcoHIV independently increased the expression of several targets, including: IL-1α [main effect of EcoHIV: F (1, 19) = 0.1221, p = 0.0479]; no main effect of B/F/TAF: F (1, 19) = 0.1221, p = 0.7306; no interaction: F (1, 19) = 3.315, p = 0.0845; **Fig. 2a**] and IL-13 [main effect of EcoHIV: F (1, 19) = 4.520, p= 0.0468; no main effect of B/F/TAF: F (1, 19) = 0.04333, p = 0.8373; no interaction: F (1, 19) = 1.096, p = 0.3083; **Fig. 2b**]. Interestingly, B/F/TAF treatment independently decreased keratinocyte chemoattractant (KC; i.e., CXCL1) expression [main effect of B/F/TAF: F (1, 19) = 6.170, p = 0.0225; no main effect of EcoHIV: F (1, 19) = 2.222, p = 0.1525; no interaction: F (1, 19) = 1.919, p = 0.1820; **Fig. 2c**].

**Figure 2.**
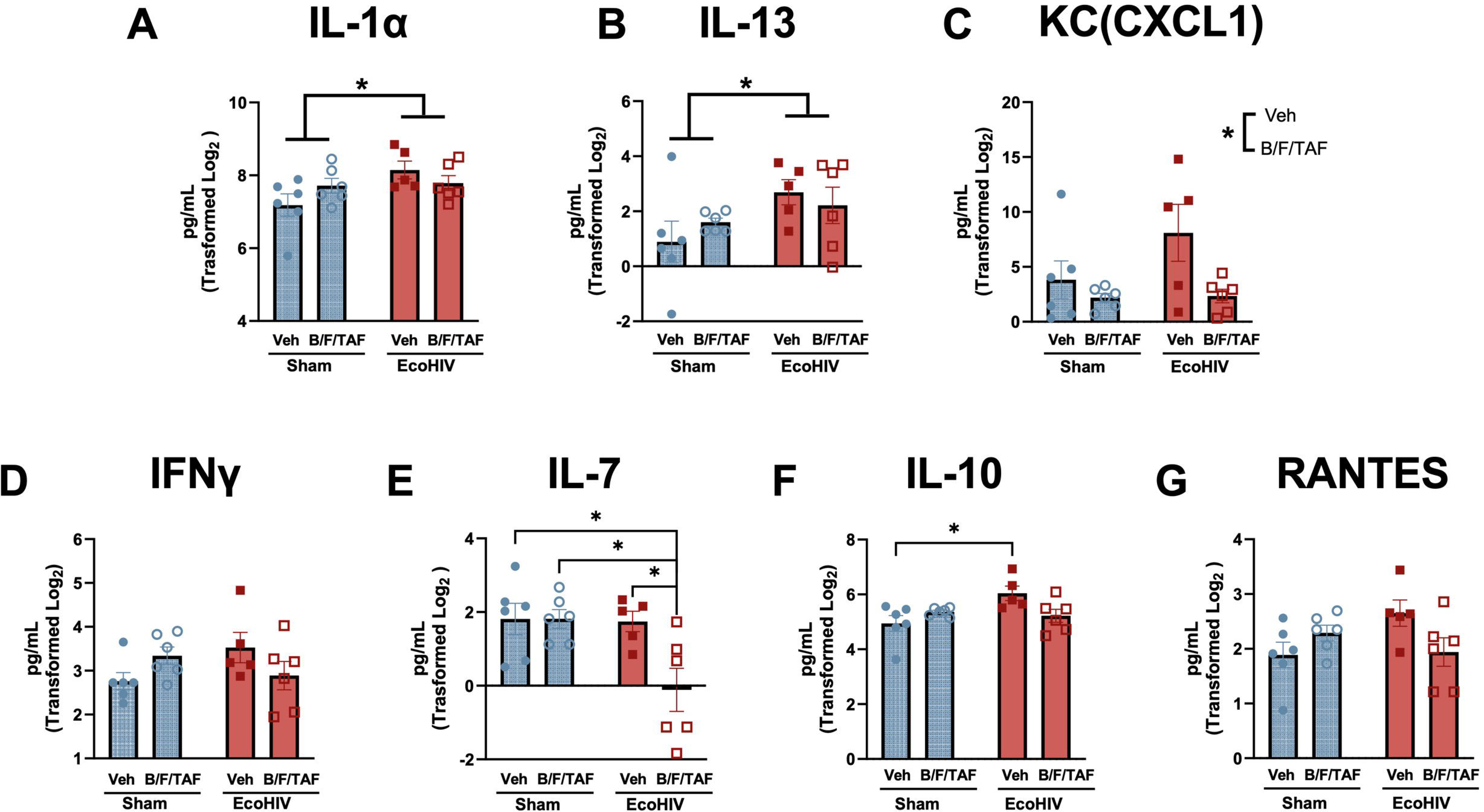
EcoHIV infection and B/F/TAF impacted immune factors in the NAc. EcoHIV, but not B/F/TAF treatment, increased expression of (**a**) IL-1α and (**b**) IL-13 in the NAc. (**c**) B/F/TAF treatment reduced expression of CXCL1/KC in the NAc independent of EcoHIV infection. EcoHIV and B/F/TAF interacted to impact (**d**) IFNɣ, (**e**) IL-7, (**f**) IL-10 and (**g**) RANTES expression levels, with selective reductions in IL-7 observed in B/F/TAF-treated mice with EcoHIV, and increased IL-10 only in vehicle-treated EcoHIV infected mice. Square and circle symbols represent sham and EcoHIV mice, close and open symbols represent vehicle and B/F/TAF treated mice, respectively. Bars represent mean +/- SEM. *p < 0.05.

Two-way ANOVAs revealed significant interactions between EcoHIV infection and B/F/TAF treatment for IFNɣ, IL-7, IL-10 and RANTES. For IFNɣ (**Fig. 2d**), a significant interaction [F (1, 19) = 5.144, p = 0.0352] but no main effects of EcoHIV [F (1, 19) = 0.01158, p = 0.9154] or B/F/TAF [F (1, 19) = 0.3436, p = 0.5647], were observed. However, *post-hoc* analyses revealed no significant comparisons. For IL-7 (**Fig. 2e**), a significant interaction [F (1, 19) = 4.917, p = 0.0390] and significant main effects of EcoHIV [F (1, 19) = 5.683, p= 0.0277] and B/F/TAF [F(1, 19) = 4.893, p = 0.0394] were observed. *Post-hoc* analysis revealed IL-7 was significantly decreased in EcoHIV mice treated with B/F/TAF compared to all other groups (*p’s < 0.05). For IL-10 (**Fig. 2f**), a significant interaction [F (1, 19) = 7.397, p= 0.0136] and main effect of EcoHIV [F (1, 19) = 4.414, p= 0.0492], but no significant main effect of B/F/TAF [F (1, 19) = 0.7504, p= 0.3972], were observed. *Post-hoc* analysis indicated that, in vehicle-treated mice, EcoHIV significantly enhanced IL-10 vs sham-inoculated mice (post-hoc: p = 0.0169). There was no difference between groups treated with B/F/TAF (p’s > 0.1). Similar to IFNɣ, analysis of RANTES expression revealed a significant interaction [F (1, 19) = 6.197, p = 0.0222] but no main effects of EcoHIV [F (1, 19) = 0.8639, p = 0.3643] or B/F/TAF [F (1, 19) = 0.4738, p = 0.4995; **Fig. 2g**] or significant *post hoc* comparisons. As NAc expression of all analytes was not normally distributed, all data were Log2 transformation prior to two-way ANOVA. The value for each analyte (uncorrected), represented as the fold-changes versus the parent control group (sham infection, vehicle treated), are visualized in the heatmap in **Supplemental Fig 1** (EcoHIV-veh: n=5; sham-B/F/TAF: n=6; EcoHIV-B/F/TAF: n=6; *p < 0.05). Values were compared by unpaired t-test, and exact p values are presented in Supplemental Table 1.

To further characterize the neuroimmune markers that were associated with EcoHIV- and B/F/TAF-treatment groups, we applied principal component analysis (PCA), a dimension reduction approach, using cytokines/chemokines that were significantly modulated in individual analyses (**Fig. 3a**). The first two principal components captured the majority of the variance (71%). Distribution along the first principal component (PC1) was driven by IL-1α, IFNɣ, IL-10 and RANTES (“Neuroimmune Cluster”), explaining 52.9% of the cumulative variance. The relationship between each marker was highly correlated. The second principal component (PC2) accounted for 19% of the variance and was positively correlated with IL-13 and negatively correlated with IL-7. The vector plot is shown in **Supplemental Fig 2**. Minimally overlapping biomarker distributions were observed between EcoHIV-infected and sham mice, with distribution along the PC2 axis corresponding with EcoHIV infection status (EcoHIV was associated with High IL-13, sham was associated with High IL-7). Distribution along PC1 corresponded with B/F/TAF treatment. Assignment to each quadrant was not equally distributed within groups of mice. In particular, 100% of vehicle-treated EcoHIV-infected mice were in the High IL-13 and High Neuroimmune quadrant. Treatment with B/F/TAF reduced the proportion of mice in this quadrant, with 33.33% of B/F/TAF-treated EcoHIV mice in this quadrant (z score two population comparison; z = 2.2887, p = 0.0220 ; **Fig. 3b**). The majority of the remaining B/F/TAF-treated EcoHIV mice (50% of total) were in the High IL-13 and Low Neuroimmune cluster. No sham-infected mice were in the High IL-13 and High Neuroimmune cluster, and only one was within the High IL-13 category. This distinct cytokine distribution indicated EcoHIV infection shifted cytokine profile toward high neuroimmune responses, and the B/F/TAF treatment attenuated this effect in EcoHIV-infected mice.

**Figure 3.**
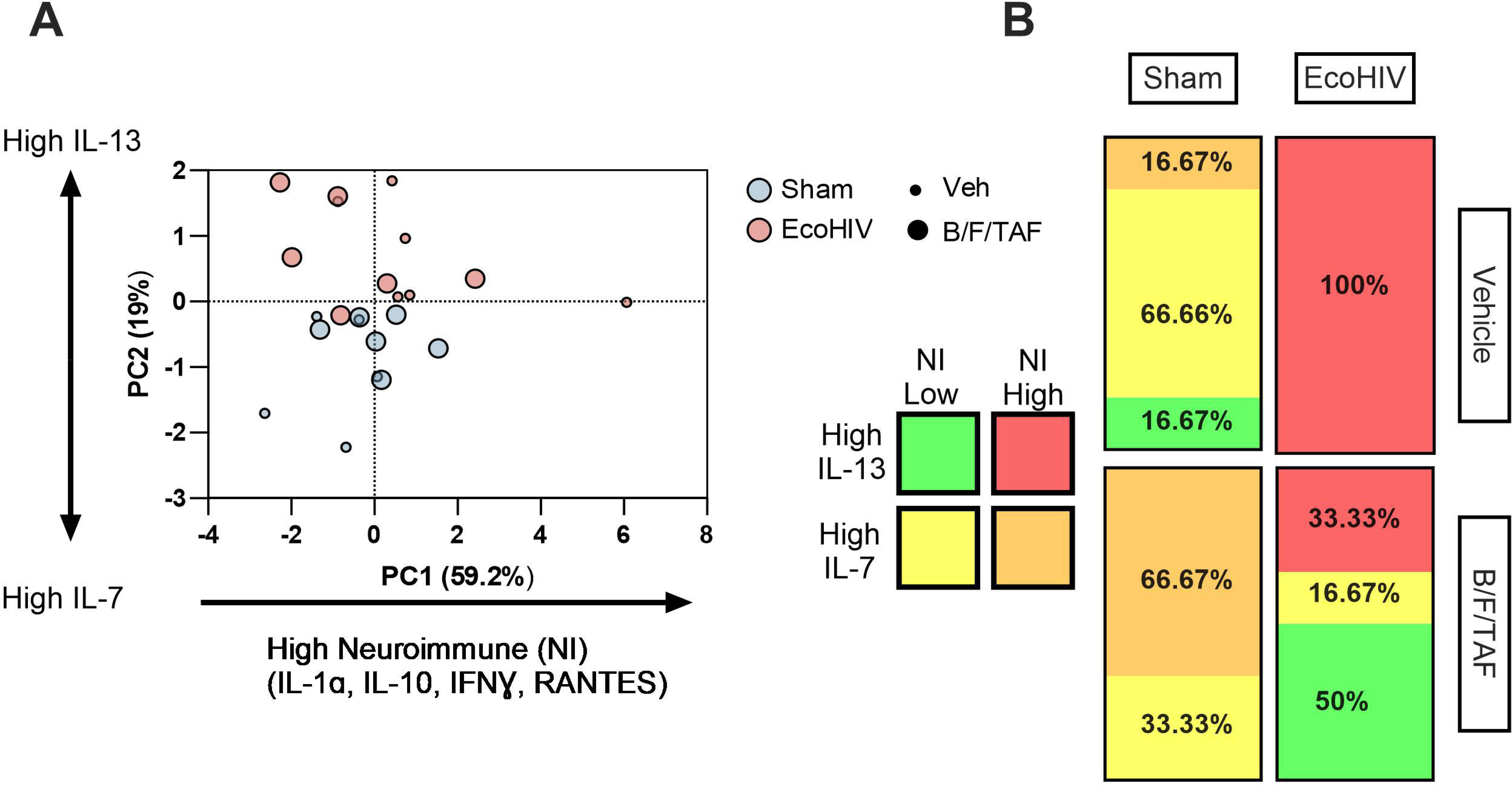
Principal component analysis (PCA) indicated distinct NAc immune profiles based on EcoHIV infection and B/F/TAF treatment. (**a**) Distribution along the first principal component (PC1) was driven by IL-1α, IFNɣ, IL-10 and RANTES (“Neuroimmune Cluster”), explaining 52.9% of the cumulative variance. The second principal component (PC2) accounted for 19% of the variance, and was positively correlated with IL-13 but negatively correlated with IL-7 expression. Sham vs EcoHIV status is indicated by color. Vehicle vs B/F/TAF treatment is indicated by symbol size. (**b**). The expression profiles of EcoHIV-infected and sham control are minimally overlapping, with distribution along the PC2 axis corresponding with EcoHIV infection status, with EcoHIV infection associated with high IL-13, and sham associated with high IL-7. Distribution along PC1 corresponded with B/F/TAF treatment, and treatment shifted immune profile to lower neuroimmune factor expression in EcoHIV infected mice.

### EcoHIV infection alters peripheral inflammatory responses in plasma

To determine peripheral inflammatory response associated with EcoHIV infection and ART, plasma was analyzed in the 32-multiplex analysis. Two cohorts of samples, including both sham- and EcoHIV-infected mice, were included in this analysis. To minimize the impact of assay batch differences, the values for each analyte were represented as the fold-change from the mean of sham-veh group within each cohort. Log_2_ transformation or square root transformation are applied to correct analytes that were not normally distributed (**Table 2**).

**Table 1.** Comparison of NAc analyte levels.

| Analyte | Effect of EcoHIV |  | Effect of B/F/TAF |  | Interaction |  |
| --- | --- | --- | --- | --- | --- | --- |
| G-CSF | F (1, 19) = 0.01514 | P=0.9034 | F (1, 19) = 0.9265 | P=0.3479 | F (1, 19) = 0.1940 | P=0.6646 |
| **IFNγ** | F (1, 19) = 0.01158 | P=0.9154 | F (1, 19) = 0.3436 | P=0.5647 | **F (1, 19) = 5.144** | **\*P=0.0352** |
| **IL-1α** | **F (1, 19) = 4.472** | **\*P=0.0479** | F (1, 19) = 0.1221 | P=0.7306 | F (1, 19) = 3.315 | P=0.0845 |
| IL-1β | F (1, 19) = 4.013 | P=0.0596 | F (1, 19) = 1.892 | P=0.1850 | F (1, 19) = 0.0001829 | P=0.9894 |
| IL-2 | F (1, 19) = 0.04298 | P=0.8380 | F (1, 19) = 0.0005031 | P=0.9823 | F (1, 19) = 0.9751 | P=0.3358 |
| IL-3 | F (1, 19) = 1.982 | P=0.1753 | F (1, 19) = 2.083 | P=0.1652 | F (1, 19) = 0.5338 | P=0.4739 |
| IL-4 | F (1, 16) = 1.172 | P=0.2951 | F (1, 16) = 2.024 | P=0.1741 | F (1, 16) = 0.7039 | P=0.4138 |
| IL-6 | F (1, 19) = 1.474 | P=0.2396 | F (1, 19) = 0.3504 | P=0.5609 | F (1, 19) = 2.145 | P=0.1594 |
| **IL-7** | **F (1, 19) = 5.683** | **\*P=0.0277** | **F (1, 19) = 4.893** | **\*P=0.0394** | **F (1, 19) = 4.917** | **\*P=0.0390** |
| IL-9 | F (1, 19) = 0.1922 | P=0.6660 | F (1, 19) = 0.7659 | P=0.3924 | F (1, 19) = 0.8635 | P=0.3644 |
| **IL-10** | **F (1, 19) = 4.414** | **\*P=0.0492** | F (1, 19) = 0.7504 | P=0.3972 | **F (1, 19) = 7.397** | **\*P=0.0136** |
| **IL-13** | **F (1, 19) = 4.520** | **\*P=0.0468** | F (1, 19) = 0.04333 | P=0.8373 | F (1, 19) = 1.096 | P=0.3083 |
| IL-12p70 | F (1, 19) = 0.002465 | P=0.9609 | F (1, 19) = 0.4221 | P=0.5237 | F (1, 19) = 0.1471 | P=0.7055 |
| IL-15 | F (1, 19) = 0.1649 | P=0.6892 | F (1, 19) = 0.3582 | P=0.5566 | F (1, 19) = 1.902 | P=0.1839 |
| IP-10 | F (1, 19) = 0.04032 | P=0.8430 | F (1, 19) = 1.502 | P=0.2353 | F (1, 19) = 0.5769 | P=0.4569 |
| **KC** | F (1, 19) = 2.222 | P=0.1525 | **F (1, 19) = 6.170** | **\*P=0.0225** | F (1, 19) = 1.919 | P=0.1820 |
| LIF | F (1, 19) = 1.954 | P=0.1783 | F (1, 19) = 0.0005130 | P=0.9822 | F (1, 19) = 0.2339 | P=0.6342 |
| LIX | F (1, 15) = 0.006022 | P=0.9392 | F (1, 15) = 0.3435 | P=0.5665 | F (1, 15) = 0.002027 | P=0.9647 |
| MCP-1 | F (1, 19) = 2.944 | P=0.1024 | F (1, 19) = 1.126 | P=0.3020 | F (1, 19) = 0.02668 | P=0.8720 |
| M-CSF | F (1, 19) = 0.1276 | P=0.7249 | F (1, 19) = 0.4704 | P=0.5011 | F (1, 19) = 1.819 | P=0.1933 |
| MIG | F (1, 19) = 0.1602 | P=0.6934 | F (1, 19) = 0.4307 | P=0.5195 | F (1, 19) = 1.887 | P=0.1855 |
| MIP-2(CXCL2) | F (1, 19) = 1.419 | P=0.2482 | F (1, 19) = 2.242 | P=0.1508 | F (1, 19) = 0.03873 | P=0.8461 |
| **RANTES** | F (1, 19) = 0.8639 | P=0.3643 | F (1, 19) = 0.4738 | P=0.4995 | **F (1, 19) = 6.197** | **\*P=0.0222** |

**Table 2.** Comparison of plasma analyte levels.

| Analyte | Effect of EcoHIV |  | Effect of B/F/TAF |  | Interaction |  |
| --- | --- | --- | --- | --- | --- | --- |
| Eotaxin | F (1, 43) = 0.05567 | P=0.8146 | F (1, 43) = 0.2570 | P=0.6148 | F (1, 43) = 0.4415 | P=0.5099 |
| G-CSF | F (1, 43) = 0.4363 | P=0.5124 | F (1, 43) = 0.02463 | P=0.8760 | F (1, 43) = 2.397 | P=0.1289 |
| IFNγ | F (1, 41) = 0.2911 | P=0.5924 | F (1, 41) = 0.2352 | P=0.6303 | F (1, 41) = 2.408 | P=0.1284 |
| IL-1α | F (1, 43) = 1.778 | P=0.1894 | F (1, 43) = 0.1580 | P=0.6930 | F (1, 43) = 0.3370 | P=0.5646 |
| IL-1β | F (1, 37) = 0.0004847 | P=0.9826 | F (1, 37) = 2.369 | P=0.1323 | F (1, 37) = 0.04638 | P=0.8307 |
| IL-2 | F (1, 31) = 2.877 | P=0.0999 | F (1, 31) = 0.02509 | P=0.8752 | F (1, 31) = 0.5275 | P=0.4731 |
| IL-3 | F (1, 42) = 1.768 | P=0.1908 | F (1, 42) = 0.02417 | P=0.8772 | F (1, 42) = 0.4318 | P=0.5147 |
| IL-4 | F (1, 43) = 0.1110 | P=0.7407 | F (1, 43) = 2.894 | P=0.0961 | F (1, 43) = 0.03413 | P=0.8543 |
| **IL-5** | F (1, 43) = 3.500 | P=0.0682 | F (1, 43) = 1.202 | P=0.2791 | **F (1, 43) = 4.080** | **\*P=0.0497** |
| **IL-6** | **F (1, 39) = 4.804** | **\*P=0.0344** | F (1, 39) = 0.8515 | P=0.3618 | F (1, 39) = 0.8436 | P=0.3640 |
| IL-9 | F (1, 43) = 0.001259 | P=0.9719 | F (1, 43) = 3.473 | P=0.0692 | F (1, 43) = 0.4027 | P=0.5290 |
| IL-10 | F (1, 43) = 0.9614 | P=0.3323 | F (1, 43) = 1.168 | P=0.2858 | F (1, 43) = 0.5482 | P=0.4631 |
| **IL-12p40** | F (1, 37) = 0.1134 | P=0.7382 | **F (1, 37) = 4.728** | **\*P=0.0361** | F (1, 37) = 0.5555 | P=0.4608 |
| IL-13 | F (1, 43) = 2.205 | P=0.1449 | F (1, 43) = 0.8116 | P=0.3727 | F (1, 43) = 0.1694 | P=0.6827 |
| IL-12p70 | F (1, 37) = 0.01623 | P=0.8993 | F (1, 37) = 0.05424 | P=0.8171 | F (1, 37) = 0.1065 | P=0.7460 |
| IL-15 | F (1, 42) = 0.8189 | P=0.3707 | F (1, 42) = 0.003764 | P=0.9514 | F (1, 42) = 0.4357 | P=0.5128 |
| IL-17 | F (1, 43) = 3.818 | P=0.0572 | F (1, 43) = 0.1825 | P=0.6713 | F (1, 43) = 0.1935 | P=0.6622 |
| IP-10 | F (1, 43) = 2.623 | P=0.1127 | F (1, 43) = 0.03397 | P=0.8546 | F (1, 43) = 0.08736 | P=0.7690 |
| KC | F (1, 43) = 0.1182 | P=0.7326 | F (1, 43) = 0.09107 | P=0.7643 | F (1, 43) = 0.4242 | P=0.5183 |
| **LIF** | **F (1, 33) = 7.941** | **\*\*P=0.0081** | F (1, 33) = 0.1217 | P=0.7294 | F (1, 33) = 0.0001909 | P=0.9891 |
| LIX | F (1, 43) = 0.1899 | P=0.6652 | F (1, 43) = 0.06386 | P=0.8017 | F (1, 43) = 0.07260 | P=0.7889 |
| MCP-1 | F (1, 40) = 0.05586 | P=0.8144 | F (1, 40) = 1.063 | P=0.3087 | F (1, 40) = 0.1222 | P=0.7285 |
| M-CSF | F (1, 43) = 1.455 | P=0.2344 | F (1, 43) = 0.9867 | P=0.3261 | F (1, 43) = 0.06422 | P=0.8012 |
| MIG | F (1, 43) = 3.117 | P=0.0846 | F (1, 43) = 0.1062 | P=0.7461 | F (1, 43) = 0.02631 | P=0.8719 |
| MIP-1α | F (1, 39) = 0.03899 | P=0.8445 | F (1, 39) = 3.739 | P=0.0605 | F (1, 39) = 0.4157 | P=0.5229 |
| MIP-1β | F (1, 43) = 0.8469 | P=0.3626 | F (1, 43) = 0.03914 | P=0.8441 | F (1, 43) = 0.6362 | P=0.4295 |
| MIP-2(CXCL2) | F (1, 32) = 0.1191 | P=0.7322 | F (1, 32) = 1.226 | P=0.2764 | F (1, 32) = 0.1690 | P=0.6837 |
| RANTES | F (1, 42) = 0.02841 | P=0.8670 | F (1, 42) = 0.3770 | P=0.5425 | F (1, 42) = 1.226 | P=0.2746 |
| TNFα | F (1, 34) = 0.2426 | P=0.6255 | F (1, 34) = 0.02013 | P=0.8880 | F (1, 34) = 0.2063 | P=0.6525 |
| VEGF | F (1, 43) = 1.382 | P=0.2462 | F (1, 43) = 0.004817 | P=0.9450 | F (1, 43) = 0.2223 | P=0.6397 |

Findings indicated that, independent of B/F/TAF treatment, EcoHIV infection reduced expression of IL-6 and LIF in plasma compared to sham mice [IL-6 (**Fig. 4a**): main effect of EcoHIV: F (1, 39) = 4.804, p = 0.0344; no main effect of B/F/TAF: F (1, 39) = 0.8515, p=0.3618; no interaction: F (1, 39) = 0.8436, p = 0.3640; LIF (**Fig. 4b**): main effect of EcoHIV: F (1, 34) = 4.768, p = 0.0360; no main effect of B/F/TAF: F (1, 34) = 0.001303, p = 0.9714; no interaction: F (1, 34) = 0.006513, p = 0.9362]. For IL-12p40 (**Fig. 4c**), a main effect of B/F/TAF treatment was observed such that IL-12p40 expression was lower in B/F/TAF treated mice [main effect of B/F/TAF: F (1, 37) = 4.728, p = 0.036; no main effect of EcoHIV: F (1, 37) = 0.1134, p = 0.7382; no interaction: F (1, 37) = 0.5555, p = 0.4608].

**Figure 4.**
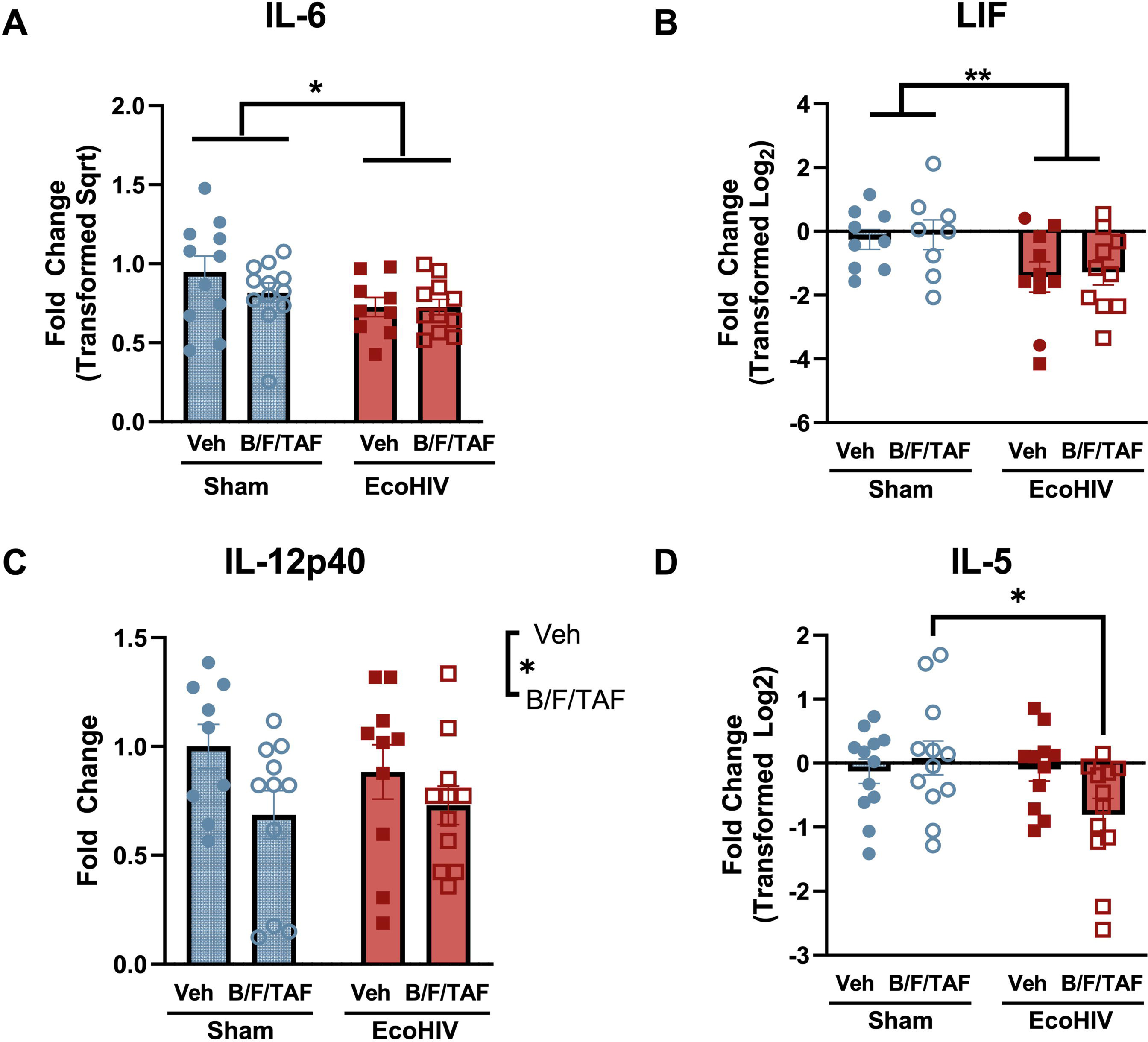
EcoHIV infection and B/F/TAF impacted immune factors in plasma. EcoHIV infection suppressed expression of (**a**) IL-6 and (**b**) LIF in the plasma independent of B/F/TAF treatment. (**c**) B/F/TAF treatment reduced IL-12p40 expression in the plasma of mice independent of EcoHIV infection. (**d**) EcoHIV and B/F/TAF interacted to alter plasma IL-5 expression such that IL-5 weas reduced only in EcoHIV-infected mice treated with B/F/TAF. Bars represent mean +/- SEM. Square and circle symbols represent sham and EcoHIV mice, closed and open symbols represent vehicle and B/F/TAF treated mice, respectively. *p < 0.05, **p < 0.01.

Expression of IL-5 (**Fig. 4d**) was driven by interactions between EcoHIV infection and B/F/TAF treatment [interaction: F (1, 43) = 4.080, p = 0.0497; no main effect of EcoHIV: F (1, 43) = 3.500, p = 0.0682; no main effect of B/F/TAF: F (1, 43) = 1.202, p = 0.2791]. *Post-hoc* analysis revealed a significant reduction in IL-5 due to B/F/TAF treatment in EcoHIV-infected mice relative to uninfected mice treated with B/F/TAF (post-hoc, p = 0.0385), but no differences were observed compared to sham-Veh and EcoHIV-Veh groups.

### EcoHIV altered the relationship between NAc Iba-1 expression and immune factor expression

To assess the effect of EcoHIV and B/F/TAF on putative microglia activation and CX3CL1 level, aliquots of the NAc lysate that were used for cytokine assays were analyzed by western blot for Iba-1 and CX3CL1 expression. Normalized Iba-1 expression was represented here as the fold change relative to the mean of sham-veh group of each cohort. No main or interaction effects of EcoHIV and B/F/TAF on NAc Iba1 expression were observed (**Fig. 5a**) [Two-way ANOVA, n= 12/group; no main effect of EcoHIV: F (1, 42) = 2.729, p = 0.1060; no main effect of B/F/TAF: F (1, 42) = 0.2158, p = 0.6446; no interaction: F (1, 42) = 1.156, p = 0.2883]. Based on our *a priori* hypothesis that EcoHIV infection would increase microglia reactivity, Iba-1 levels between sham-Veh and EcoHIV-Veh groups were compared. An unpaired t-test demonstrated a significant increase of Iba-1 expression in EcoHIV-infected, vehicle-treated mice compared to sham, vehicle treated mice [t (21)= 2.168, p = 0.0418, 95% C.I. = 0.01390 to 0.6687; **Fig. 5d**] indicating that 5 weeks of EcoHIV infection significantly upregulated Iba-1 expression in the NAc. No effects of EcoHIV or B/F/TAF treatment were observed on CX3CL1 expression in the NAc [EcoHIV: F (1, 42) = 0.0007631, p = 0.9781; B/F/TAF: F (1, 42) = 0.05451, p = 0.8165; interaction: F (1, 42) = 0.09385, p = 0.7609; **Fig. 5b**]. No differences were observed in CX3CL1 expression between sham-Veh and EcoHIV-Veh mice [t (21) =0.1846, p = 0.8553, 95% C.I. = -0.3007 to 0.2517; **Fig. 5d**]. Representative blots are shown in **Fig 5c** (full blots in **Supplemental Figure 4**).

**Figure 5.**
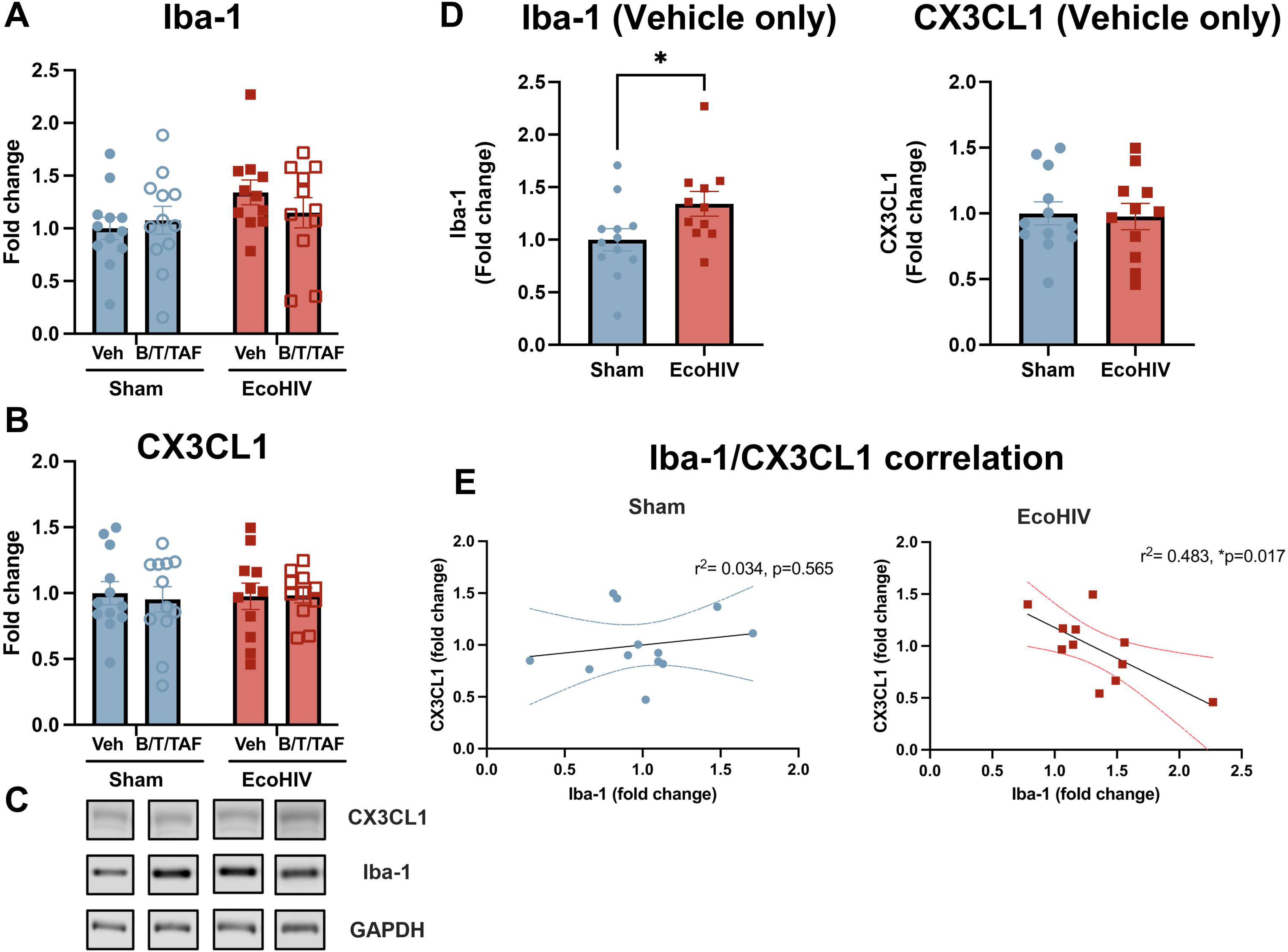
EcoHIV infection determined the relationship between Iba-1 and NAc expression of immune factors. When compared by ANOVA, no effect of EcoHIV infection or B/F/TAF was observed on NAc (**a**) Iba-1 expression or (**b**) CX3CL1 expression. (**c**) Representative Iba-1, CX3CL1 and internal control GAPDH western blots. Based on an *a priori* hypotheses, overall mean NAc expression of Iba-1 and CX3CL1 were compared between Sham and EcoHIV-infected mice. EcoHIV infected mice exhibited higher levels of Iba-1, but not CX3CL1 (**d**). There was no relationship between Iba-1 and CX3CL1 expression in sham-infected mice, while expression was negatively correlated in EcoHIV infected mice (**e**). *p < 0.05.

Previous studies have shown that microglia activation is negatively associated with fractalkine system expression. Thus, we performed simple linear regression analysis between Iba-1 and CX3CL1 in sham and EcoHIV vehicle treated groups. Linear regression showed a negative correlation with Iba-1 and CX3CL1 levels in EcoHIV-Veh mice (R^2^= 0.4825, p = 0.0177); whereas no correlation was observed between Iba-1 and CX3CL1 in sham-Veh group (R^2^= 0.03419, p = 0.5651; **Fig. 5e**). This result may indicate that EcoHIV-induced increases in Iba1 expression are at least partially suppressed by CX3CL1 signaling within the NAc.

To determine whether Iba-1 expression was associated with NAc expression of immune factors, simple linear regression analysis was performed (**Supplemental Fig 3**). We observed differential patterns of relationships in particular in EcoHIV infected mice that were not treated with B/F/TAF, especially the inversion of the relationship between CX3CL1 expression and Iba-1 and LIF and Iba-1. These correlations are summarized in **Table 3**.

**Table 3.** Correlations between Iba-1 expression and NAc analytes.

|  | Sham-Veh |  | EcoHIV-Veh |  | Sham-B/F/TAF |  | EcoHIV-B/F/TAF |  |
| --- | --- | --- | --- | --- | --- | --- | --- | --- |
|  | Pearson's r | p value | Pearson's r | p value | Pearson's r | p value | Pearson's r | p value |
| M-CSF | -0.068222718 | 0.8978 | 0.839266527 | #0.0755 | 0.117159662 | 0.8251 | -0.206332678 | 0.6949 |
| MIG | -0.44952674 | 0.3711 | 0.817740051 | #0.0908 | 0.389758049 | 0.445 | 0.824767601 | **\*0.0434** |
| LIF | -0.177022696 | 0.7372 | 0.745987306 | 0.1477 | -0.512975355 | 0.298 | -0.682293114 | 0.1354 |
| IL-4 | -0.579415488 | 0.3059 | 0.728241495 | 0.163 | -0.074996179 | 0.8877 | -0.892826859 | 0.1072 |
| KC | -0.026069555 | 0.9609 | 0.589752177 | 0.2952 | -0.010616142 | 0.9841 | -0.118709927 | 0.8228 |
| MCP-1 | -0.412394917 | 0.4165 | 0.375521109 | 0.5334 | -0.894373728 | **\*0.0161** | -0.928418899 | **\*\*0.0075** |
| G-CSF | 0.206900311 | 0.6941 | 0.35096456 | 0.5625 | -0.426215677 | 0.3994 | -0.403295956 | 0.4279 |
| IL-13 | 0.511634468 | 0.2995 | 0.225110668 | 0.7158 | -0.086223279 | 0.871 | 0.736262156 | 0.0952 |
| IL-2 | 0.404048716 | 0.4269 | 0.122146027 | 0.8449 | -0.379897186 | 0.4576 | -0.091448348 | 0.8632 |
| IL-7 | -0.390715732 | 0.4437 | 0.117940131 | 0.8502 | -0.47748701 | 0.3382 | 0.054385453 | 0.9185 |
| IL-3 | -0.00745248 | 0.9888 | -0.166397688 | 0.7891 | -0.117165655 | 0.8251 | -0.38481192 | 0.4513 |
| IL-6 | 0.458129801 | 0.3609 | -0.220218491 | 0.7219 | 0.474829516 | 0.3413 | 0.511465649 | 0.2997 |
| IL-9 | -0.867881407 | **\*0.025** | -0.285552826 | 0.6414 | -0.884342292 | **\*0.0193** | -0.833558575 | **\*0.0392** |
| RANTES | 0.203542734 | 0.6989 | -0.400252342 | 0.5043 | -0.277697902 | 0.5942 | -0.787683252 | #0.0628 |
| IL-15 | -0.420596675 | 0.4063 | -0.407445935 | 0.496 | -0.620689799 | 0.1885 | -0.629115677 | 0.1808 |
| IFNγ | -0.201985846 | 0.7011 | -0.461319212 | 0.4342 | -0.786791051 | #0.0633 | -0.80836188 | #0.0516 |
| IP-10 | 0.210078974 | 0.6895 | -0.59978229 | 0.285 | -0.339432646 | 0.5104 | -0.783924787 | #0.065 |
| IL-10 | 0.611275269 | 0.1973 | -0.607840565 | 0.2768 | -0.060485338 | 0.9094 | -0.728361676 | 0.1007 |
| IL-1α | 0.398494481 | 0.4339 | -0.647407914 | 0.2376 | -0.587864175 | 0.2198 | -0.955432251 | **\*\*0.0029** |
| IL-12p70 | -0.746521652 | #0.0882 | -0.689806805 | 0.1975 | -0.937797428 | **\*\*0.0057** | -0.91026093 | **\*0.0117** |
| MIP-2 | -0.472472412 | 0.344 | -0.741169537 | 0.1518 | 0.7911121 | #0.0609 | 0.294744877 | 0.5707 |
| IL-1β | 0.074942205 | 0.8878 | -0.8025595 | 0.1021 | -0.401488171 | 0.4301 | -0.564794703 | 0.2429 |
| CX3CL1 | 0.796233045 | #0.0581 | -0.959630735 | **\*\*0.0097** | 0.712478158 | 0.1121 | 0.58665987 | 0.221 |

## Discussion

The current findings demonstrate independent and interactive effects of EcoHIV infection and ART exposure on peripheral and brain immune response in mice. Given previous findings indicating sensitivity of brain reward circuitry to perturbation in models of HIV infection, these experiments focused on dysregulation within the NAc, where increased expression of IL-1α and IL-13 were induced by EcoHIV infection. Within the NAc, treatment with the ART combination B/F/TAF independently reduced expression of CXCL1/keratinocyte-derived chemokine (KC) and interacted with EcoHIV infection to alter expression of IL-7, IL-10, IFNɣ and RANTES. Changes in chemokine expression in the NAc were not accompanied by gross alterations in Iba-1 expression, a putative marker of microglia, though the relationship between Iba-1 and chemokine expression was determined by EcoHIV and B/F/TAF treatment status. Notably, the immune targets for which alterations were observed in plasma were distinct from those observed within the NAc. In plasma, EcoHIV infection reduced expression of IL-6 and leukemia inhibitory factor (LIF), a member of the IL-6 family. Treatment with B/F/TAF reduced expression of IL-12p40 and interacted with EcoHIV infection to selectively reduce IL-5 expression in EcoHIV-infected mice.

Neuroimmune response is an important CNS outcome following HIV infection, which may have deleterious effects on neural function through protracted inflammation. Neuroimmune dysregulation within the CNS can result from infiltration of HIV-1 infected macrophages across the blood brain barrier and subsequent infection of resident CNS immune cells that maintain persistent CNS viral reservoirs (Crowe et al., 2003; Veenstra et al., 2017b, 2017a). HIV-1 infected macrophages and microglia release viral proteins and inflammatory cytokines/chemokines, which in turn dysregulates immune signaling and disrupts homeostatic neuronal function (Fitting et al., 2013; Williams et al., 2014; Jones et al., 2016). It has been well-documented that HIV infection is associated with interleukin-1 (IL-1) family regulation in the CNS (Merrill and Chen, 1991; Flora et al., 2005; Brabers and Nottet, 2006; McLaurin et al., 2021). A large majority studies on HIV infection and the IL-1 family have focused on the IL-1β and demonstrate that IL-1β is a major mediator of HIV-associated neuroinflammation (Gemma et al., 2000). For example, in rats treated with HIV-1 gp120, IL-1β upregulation within the cortex is associated with neuronal dysfunction and cognitive deficits (Festa et al., 2015). While IL-1α is a key pro-inflammatory cytokine in the IL-1 family, a clear role for IL-1α has not been identified in preclinical HIV models. IL-1α release is upregulated in response to brain injuries via activated microglia, which likely contributes to chronic inflammation and neurocognitive impairment (Brough and Denes, 2015). Others have reported upregulated IL-1α in models of co-occurring HIV (Tat protein model) and drug exposure (Theodore et al., 2006), and further that elevation of IL-1α in the cerebrospinal fluid of PLWH is positively associated with risk for cognitive impairment (Williams et al., 2021). We found that EcoHIV-infected mice exhibit upregulated IL-1α in the NAc despite ART treatment, suggesting that persistent CNS levels of IL-1α may be present even in virally-suppressed PLWH.

EcoHIV infection also increased IL-13 in the NAc. The function of IL-13 is highly context dependent. For example, IL-13 is postulated to modulate neuronal homeostasis in healthy conditions, and it is upregulated following brain injury (Mori et al., 2017). In multiple brain injury models, IL-13 upregulation had a neuroprotective effect in multiple brain injury models (Mori et al., 2017; Kolosowska et al., 2019; Li et al., 2023). Further, IL-13/IL-13 receptor alpha 1(IL-13R1) interacted prevent dopaminergic neuronal loss in a model of chronic restraint stress (Mori et al., 2017). Peripheral IL-13 administration also appears to be neuroprotective, such as by inducing anti-inflammatory, microglia-mediated immune responses that exhibit neuroprotective effects in a mouse model of ischemic stroke (Kolosowska et al., 2019). Further, IL-13 upregulation promotes phosphorylation of NMDA and AMPA glutamate receptors that can drive synaptic plasticity. While this may protect neurons from excitotoxic death following brain injury (Li et al., 2023), this effect on glutamate receptor expression could produce cognitive impairments akin to those observed in HAND (O’Donnell et al., 2006). Increased IL-13 release from reactive microglia has also been shown to be neurotoxic following lipopolysaccharide injection. In this model, increased IL-13 immunoreactivity was further associated with astrocyte damage and blood brain barrier (BBB) disruption (Hong et al., 2022). Thus, it is essential to consider the function of IL-13 in the context of specific disorders and disease states, including the state of disease progression, and the specific function of increased NAc IL-13 in the EcoHIV model remains a promising area for future study.

Elevated IL-1α and IL-13 expression in the NAc - a key neural substrates of reward learning and processing - of EcoHIV-infected mice may be associated with excitatory neuronal adaption. IL-1 released from reactive microglia in drug addiction model is involved in enhancing glutamate transmission through downregulate glutamate transporter (GLT-1) (Prow and Irani, 2008) and potentiates the activation of ionotropic glutamate receptors (e.g., AMPA, NMDA) in the NAc (Viviani et al., 2003; Liu et al., 2013). IL-1 and IL-13 effects on synaptic plasticity are thought to be associated with long-term potentiation to the excitatory glutamatergic neurons (Viviani et al., 2003; Khairova et al., 2009; Gipson et al., 2021; Li et al., 2023), which is overlapping with the action of addictive substances in the CNS. Thus, exposure of neurons to high IL-1α and IL-13 may induce differential neuronal susceptibility to excitatory glutamate stimulation. This may suggest one mechanism by which PLWH are at greater risk for development of SUD when exposed to addictive drugs. This finding is informative to the research focus on the HIV-associated comorbidities, including SUD, as the number of PLWH with SUD continues to increase (Buch et al., 2011; Giacometti and Barker, 2019).

Interestingly, a selective reduction in NAc IL-7 was observed in B/F/TAF-treated mice infected with EcoHIV. IL-7 effects are mediated through binding to the receptor IL-7R. IL-7/IL-7R signaling is a critical regulator of CD4+ T cell homeostasis (Vandergeeten et al., 2013). HIV infection can disrupt intracellular signaling downstream of IL-7R by preventing STAT5 nuclear localization. This may contribute to subsequent loss of CD4s, which could thus worsen immune activation (Catalfamo et al., 2008; Landires et al., 2011). Elevated levels of serum or plasma IL-7 are observed in PLWH and animal models (Napolitano et al., 2001; Camargo et al., 2009), and IL-7 administration promotes HIV viral persistence during ART treatment by promoting rapid viral production (Vandergeeten et al., 2013). One potential mechanism that may contribute to ART-mediated rescue of CD4+ and CD8+ T cells counts is through enhanced IL-7 clearance via increased receptor availability of IL-7R on T cells (Hodge et al., 2011). Altogether, these findings point toward a hypothesis that increases in IL-7 and dysregulation of IL-7/IL-7R signaling may contribute to the pathophysiology of HIV, and that reduced IL-7 may help mediate the therapeutic efficacy of ART. In the present study, IL-7 was only decreased in EcoHIV-infected mice with B/F/TAF treatment. While the current study did not assess CNS EcoHIV expression, it is possible that B/F/TAF-mediated suppression of IL-7 within the NAc may reduce CNS infection, which would align with evidence of ART-mediated reductions in IL-7 may reduce HIV reservoirs (Hodge et al., 2011; Goonetilleke et al., 2019).

In addition to IL-7, interactions between EcoHIV infection and B/F/TAF were observed in IFNɣ, IL-10 and RANTES, which are associated with neuropathogenesis and persistence of HIV-associated neurocognitive deficits (Monteiro et al., 2016; Porro et al., 2020; Williams et al., 2021). Specifically, NAc IL-10 expression was significantly greater in vehicle-treated EcoHIV-infected mice to Sham-Veh controls. Upregulated IL-10 also has been reported to play a role in viral load persistence and T cell impairment in the context of HIV (Planès et al., 2018; Harper et al., 2022). While post-hoc analyses were nonsignificant, parallel group trends and significant interaction effects were observed for both IFNɣ and RANTES, consistent with EcoHIV-infection promoting neuroimmune response and a role of ART in attenuating neuroimmune response in the EcoHIV model. This aligns with our PCA analysis, as vehicle-treated EcoHIV mice were distributed in the quadrant associated with High IL-13 and High Neuroimmune response. Treatment with B/F/TAF shifted the profile of EcoHIV-infected mice to a lower Neuroimmune response, without changing IL-13 levels. Notably, no sham-infected animals were in the High-IL-13/High Neuroimmune response quadrant. Further, B/F/TAF treatment in sham mice did not have the same effects on immune profile, consistent with independent and interactive effects of ART treatment on immune status in the CNS.

Plasma immune profile is widely used as a biomarker for monitoring HIV disease progression. Peripheral immune activation in response to HIV is essential for protective immunity against viral invasion. In general, plasma cytokine dysregulation during acute and chronic HIV infection is associated with CD4+ and CD8+T cell activation status, viral load set-point, and efficacy of ART therapy (Shete et al., 2020; Ngcobo et al., 2022). Notably, in the current study, the observed changes in immune profiles were not congruent between plasma and the NAc. In EcoHIV-infected mice, downregulated plasma IL-6 and LIF expression was observed regardless of B/F/TAF treatment. IL-6 is widely documented as a proinflammatory cytokine that is elevated by HIV-infected monocytes and macrophages (Breen et al., 1990). Increased circulating IL-6 level is linked to HIV replication, decreased CD4+ cell counts and elevated inflammatory response in PLWH (Borges et al., 2015). Blockade of IL-6 activity has been reported to attenuate these effects (Rodriguez et al., 2020), indicating IL-6 level can serve as both a marker for HIV inflammation outcome and play a role in reversing these outcomes. Leukemia inhibitor factor (LIF) is a chemokine that is associated with anti-inflammatory and T cell protection properties in HIV-1 infection (Tjernlund et al., 2006). It has been indicated LIF can act as HIV-1 suppressor (Patterson et al., 2001; Tjernlund et al., 2007). LIF inhibits HIV-1 replication via STAT3 signaling (Tjernlund et al., 2007), and the loss of LIF may contribute to HIV transmission and increased viral load (Patterson et al., 2001). We also observed a reduction in plasma IL-12p40 due to B/F/TAF treatment. IL-12p40 is a proinflammatory cytokine known to promote autoimmune inflammation(Gee et al., 2009), which occurred independent of EcoHIV infection status. These findings are consistent with the interpretation that B/F/TAF treatment is not only interacting with EcoHIV infection to alter outcomes, as observed for IL-7 in the NAc or IL-5 in the periphery, but - similar to CXCL1/KC findings within the NAc - is independently altering immune targets even in the absence of infection.

Within the CNS, HIV mRNA is thought to be predominantly expressed in microglia. High co-localization between Iba1 and HIV-1 mRNA has been observed in the frontal-striatal area in HIV-1 Tg rats model (Li et al., 2021). In the EcoHIV model following an intracranial or retroorbital route of infection – a different route than the peripheral inoculation used in the current study – EcoHIV-EGFP signal was highly co-localized with Iba-1 in the CNS (Li et al., 2021). In addition, intracranial EcoHIV infection induced reactive morphology in microglia, increased expression of inflammatory factors, and neuronal and synaptic impairments (but not apoptosis) in the basal ganglia (Kelschenbach et al., 2019). These results align with our findings that following 5 weeks of EcoHIV infection, independent of B/F/TAF treatment, immune activation in the NAc was increased (including IL-1α, IL-10 and IL-13), potentially driven by activated microglia. This suggests that EcoHIV may facilitate microglia activation and associated immune responses in the NAc as early as 5 weeks post-infection. Although the intracerebral EcoHIV injection model showed a drastic change in Iba-1 expression and immune response in the brain, a caveat for this model is that this injection procedure and large amount of viral load within the CNS can cause mechanical damage and viral protein-induced neurotoxicity, which may be independent of viral infection *per se*. The findings in the current manuscript indicate more subtle changes in the NAc due to peripheral EcoHIV infection, which may recapitulate the mild, low-grade neuroimmune activation that characterizes neuroHIV and HAND in PLWH (Davis et al., 1992; Valcour et al., 2012; Gaskill et al., 2017).

The CX3CL1-CX3CR1 signaling axis mediates microglial and neuronal cell communication in pathological conditions including HIV (Tong et al., 2000; Pereira et al., 2001). The interaction between neurons that release CX3CL1 and microglia expressing CX3CR1 acts to attenuate microglia activation and promote intracellular signaling pathways for neuroprotection responses. Early studies demonstrated CX3CL1-CX3CR1 signaling inhibits excitatory postsynaptic glutamatergic transmission that protect neurons from excitotoxic damage (Chapman et al., 2000; Cardona et al., 2006). CX3CL1 can exert anti-inflammatory effects; it suppresses the release of nitric oxide (NO), IL-6, and tumor necrosis factor (TNF)-α by activated microglia in a dose-dependent manner (Mizuno et al., 2003). CX3CL1 has also been reported to protect hippocampal neurons from HIV toxic protein gp120 damage, and this effect is blocked by CX3CR1 blockage or CX3CR1 knockout (Meucci et al., 2000; Ru et al., 2019), suggesting a critical contribution of CX3CL1-CX3CR1 signaling in neuroprotection in the context of neuroHIV. While CX3CL1 appears to be neuroprotective in some circumstances, enhanced CX3CL1-CX3CR1 signaling is also involved in neuropathological conditions. For example, increased expression of CX3CL1 and CX3CR1 is associated with abnormal microglial activation and neuronal apoptosis in epilepsy models (Xu et al., 2012). CX3CL1 upregulation was also reported in the brain of PLWH with cognitive impairment without on ART therapy (Pereira et al., 2001), indicating the disruption of the CX3CL1-CX3CR1 axis by HIV may have either beneficial or detrimental effects on CNS depending on disease state. We hypothesized that EcoHIV infection would dysregulate NAc CX3CL1 expression. However, no significant difference in mean CX3CL1 levels was observed between sham and EcoHIV-infected mice. Though not assessed in the current study, EcoHIV infection may have impacted CX3CR1 expression levels. CX3CL1 release is activity-dependent and inflammatory stimuli drive CX3CL1 release, which consequently regulates microglia activation and exerts neuroprotective effects (Cardona et al., 2006; Durán Laforet Dorothy Schafer and Dorothy Schafer, 2024). Indeed, Iba-1 and CX3CL1 were inversely correlated in EcoHIV-infected mice, such that higher levels of Iba-1 were associated with reduced CX3CL1. This relationship was absent in sham mice. The directionality of this relationship in EcoHIV mice is not clear from the current study. One possibility is that in EcoHIV, increased CX3CL1-CX3CR1 signaling suppresses microglia activation in the NAc. This finding is in agreement with previous studies that microglia activation is negatively regulated by the CX3CL1-CX3CR1 system (Cardona et al., 2006; Xie et al., 2013).

Importantly, the EcoHIV model does not fully recapitulate HIV-1 infection. One notable difference includes the absence of gp120, which is a known mediator of deleterious immune and neural outcomes in HIV-1 infection (Gemma et al., 2000; Hill et al., 2019). While consistent with general peripheral and CNS immune system dysregulation observed in rodent models of HIV (Fitting et al., 2013; Melendez et al., 2016; Namba et al., 2023a), the time course of EcoHIV infection differs from HIV-1 (Gu et al., 2018). In the current study, we report that EcoHIV-infected mice showed viral DNA burdens of up to 0.3-5 x 10^3^ viral DNA copies per 10^6^ spleen cells with 5 weeks of infection. Treatment with B/F/TAF starting one week following initial infection did not alter EcoHIV viral DNA burden. ART treatment can safely and effectively suppress viraemia to an undetectable level. However, ART does not eliminate HIV DNA (Simonetti and Kearney, 2015; NIAID, 2018). EcoHIV appears to establish and maintain low DNA burden in spleen cells. Gu and colleagues examined EcoHIV viremia by measuring viral RNA in blood throughout 8 weeks of infection and showed that EcoHIV infection achieves peak viral load within the first week of infection. Peripheral EcoHIV RNA copies showed a significant decline by week 3 post-infection and dropped to undetectable level at week 8. In contrast, viral DNA levels plateau in the spleen following a transient peak after inoculation, and the DNA burden can remain stable for 15 months after infection (Gu et al., 2018). This suggests that ART is not necessary for the suppression of viral load within the periphery of wild type mice, indicating an endogenous antiviral immune response that limits EcoHIV replication in mice. Once EcoHIV establishes a chronic infection, the virus persists at a low burden in spleen cells in the form of latent viral reservoir and ART is not sufficient to restrict EcoHIV viral burden. Importantly, it is increasingly apparent that alterations in neurocognitive function do not require persistent elevations in viral load. For example, in a model assessing the contribution of peripheral macrophages migrating to the brain within weeks following systemic EcoHIV infection, ART treatment following infection was unable to attenuate either viral burden or cognitive impairment (Gu et al., 2018). Similarly, mild cognitive impairment has been observed in PLWH with controlled HIV load (Peterson et al., 2014). Together, these findings identify lasting neuroimmune consequences of EcoHIV infection beyond the known windows of high circulating viral loads (Potash et al., 2005), which are modulated by, but not reversed in the presence of, ART. This represents one potential mechanism by which neurocognitive impairments may develop in the virally suppressed state.

## Conclusions

Together, findings in these studies identify interactive and independent effects of both EcoHIV infection and ART treatment on expression of immune targets in the periphery and within the NAc. This underscores the need for preclinical models to emphasize inclusion of ART as a variable in investigating both systemic and CNS inflammatory outcomes and the potential for ART to alter neuroimmune function.

## Acknowledgements

This research was supported by NIH awards DP2DA051907 (JMB), R03DA047919 (JMB), R01AG081929 (JGJ) and R21DA056309 (MPI Jackson, Klase, Meucci), and pilot awards from The Comprehensive Neuro-AIDS Center Grant P30MH092177-9 (JMB and MDN). The EcoHIV-NDK plasmid was a gift from David Volsky.

