## Supplemental Figures for "Effects of antiretroviral treatment on central and peripheral immune response in mice with EcoHIV infection"

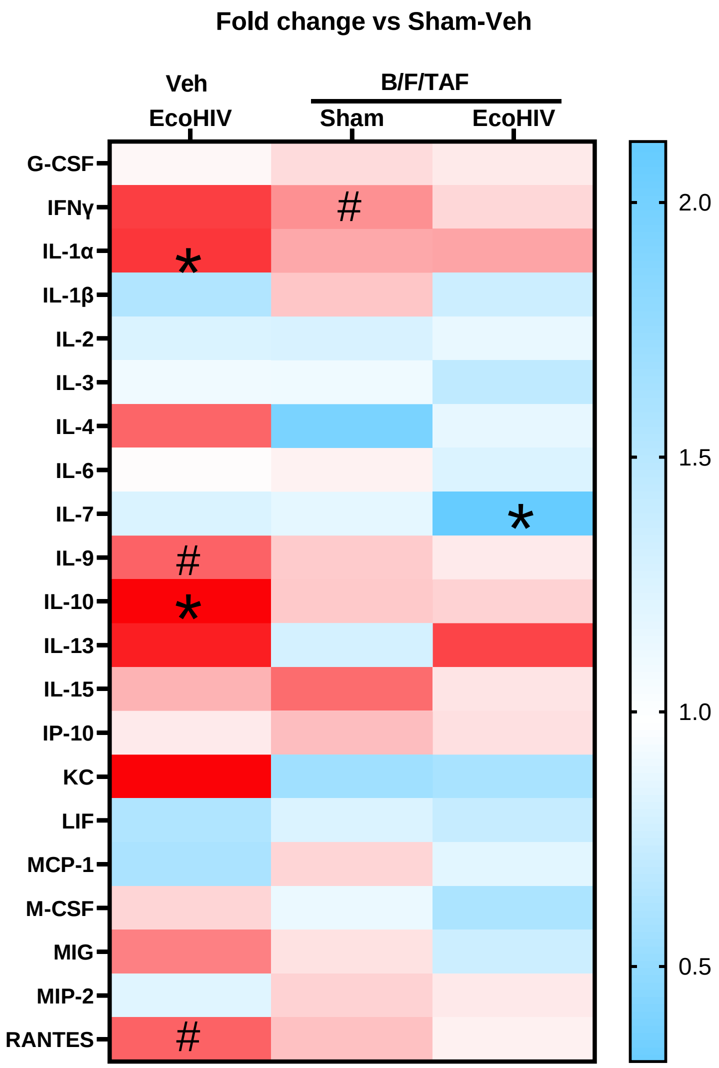


**Supplemental Figure 1. NAc expression of inflammatory factors versus vehicle control.** Expression of NAc chemokine and cytokines in EcoHIV-Veh, sham-B/F/TAF and EcoHIV-B/F/TAF treated mice represented as the fold-change verse sham-Veh group shown as heatmap (fold increase indicated in red, decrease in blue). *p < 0.05, # p<0.1

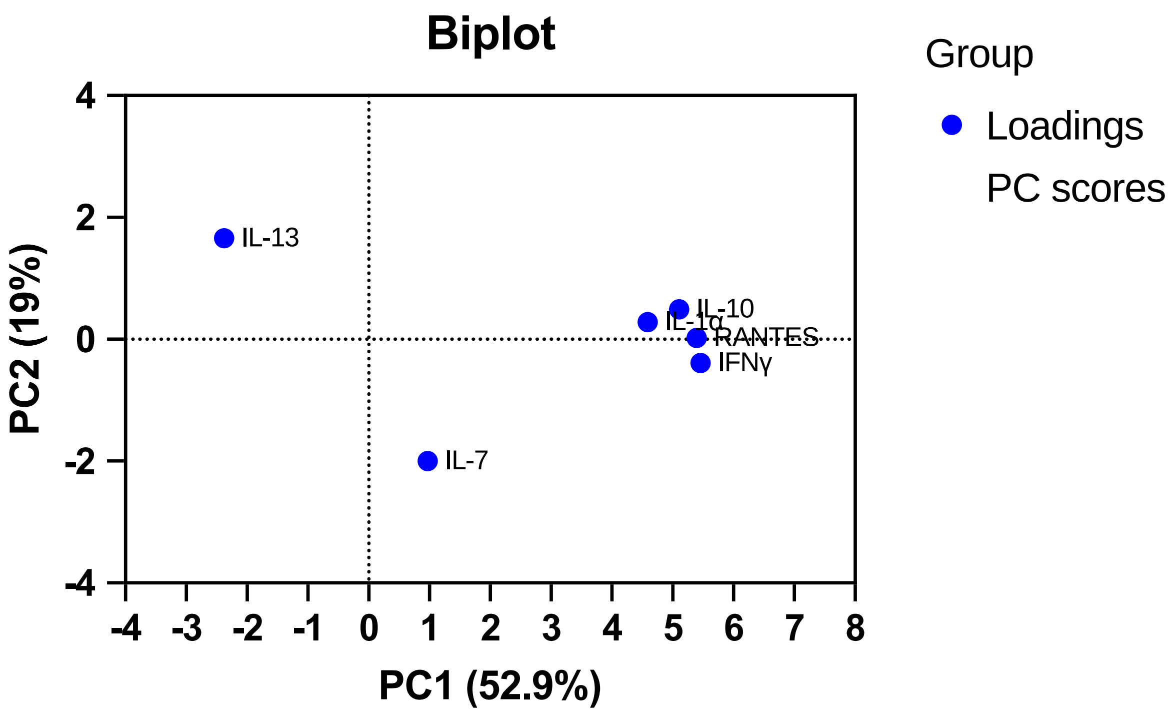


**Supplemental Figure 2. Biplot of the attributes of PCA.** The biplot shows the impact of each biomarker on each of PCs. IL-1α, IFNɣ, IL-10 and RANTES contribute to PC1 loading and are positively correlated (clustered). IL-13 and IL-7 contribute to PC2 loading and they are negatively correlated (opposite sides of the biplot origin).

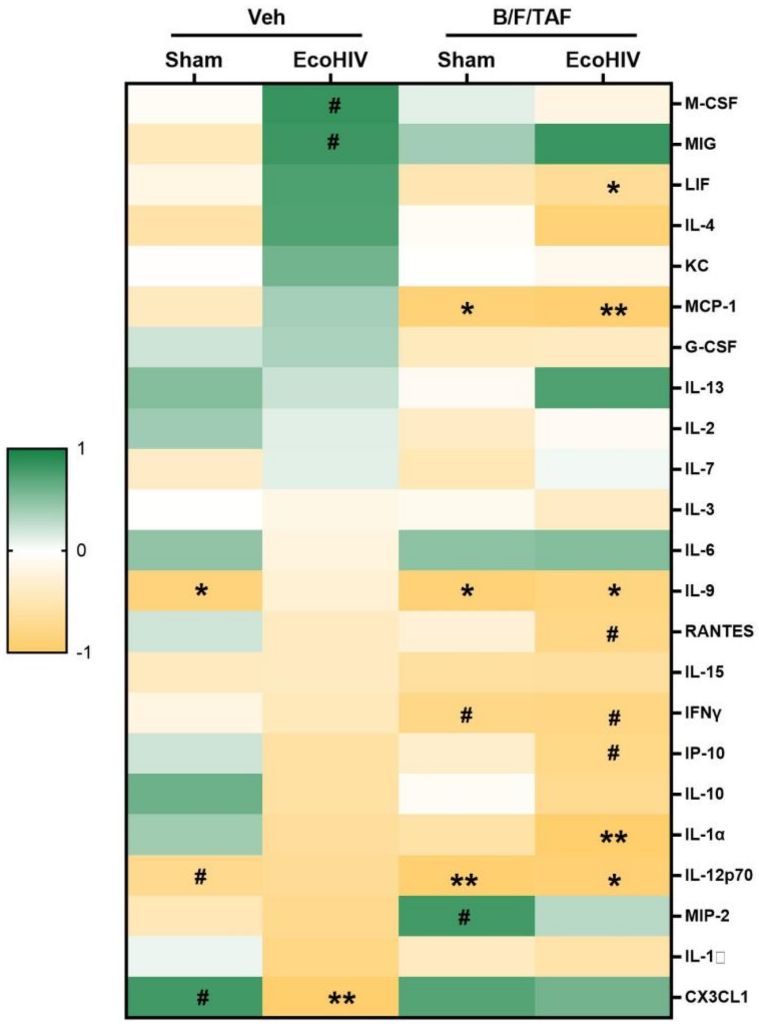


**Supplemental Figure 3.** The relationship between Iba-1 expression and expression of immune factors in the NAc was modulated by EcoHIV infection and B/F/TAF-treatment status, with distinct patterns of relationships observed in particular in the EcoHIV-infected, vehicle-treated mice. Colors represent $r^2$ values of correlations (green is positive, yellow is negative). *$p < 0.05$, **$p < 0.01$, # $p<0.1$.

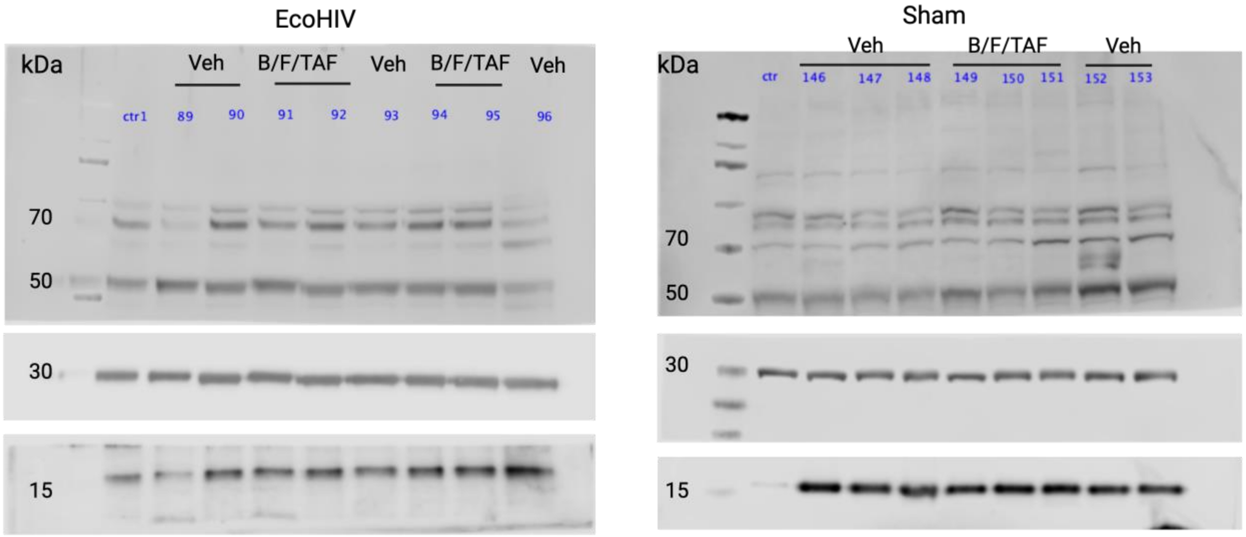


**Supplemental Figure 4.** The complete western whole blot for CX3CL1, Iba-1 and GAPDH.
